# Rapid detection of G6PD deficiency SNPs using a novel amplicon-based MinION Sequencing Assay

**DOI:** 10.1101/2025.06.23.661218

**Authors:** Costanza Tacoli, Mariana Kleinecke, Kian Soon Hoon, Hidayat Trimarsanto, Muhammad Nadeem, Rotha Eam, Nimol Khim, Angela Rumaseb, Benedikt Ley, Dean Sayre, Sochea Phok, Dysoley Lek, Benoit Witkowski, Jimee Hwang, Sarah Auburn, Ivo Mueller, Ric N Price, Jean Popovici

**Affiliations:** Malaria Research Unit Institut Pasteur du Cambodge, Phnom Penh, Cambodia; Walter and Eliza Hall Institute, Melbourne, Australia; Global and Tropical Health Division, Menzies School of Health Research, Charles Darwin University, Darwin, Northern Territory, Australia; Eijkman Research Center for Molecular Biology, National Research and Innovation Agency, Jakarta, Indonesia; U.S. President’s Malaria Initiative, Malaria Branch, Centers for Disease Control and Prevention, Atlanta, USA; PMI Impact Malaria, Population Services International, Phnom Penh, Cambodia; National Center for Parasitology, Entomology and Malaria Control, Phnom Penh, Cambodia; Mahidol-Oxford Tropical Medicine Research Unit, Mahidol University, Bangkok, Thailand; Centre for Tropical Medicine and Global Health, Nuffield Department of Medicine, University of Oxford, Oxford, UK; University of Melbourne, Melbourne, Australia; Infectious Disease Epidemiology and Analytics G5 Unit, Institut Pasteur, Paris, France

**Author notes:** equal contribution (Costanza Tacoli, Mariana Kleinecke). Corresponding author: Jean Popovici.

## Abstract

*Plasmodium vivax* malaria remains a significant global health challenge, complicated by the parasite’s ability to form dormant liver stages (hypnozoites) that cause relapses. Radical cure of *P. vivax* malaria requires administration of a hypnozoitocidal drug, such as primaquine or tafenoquine. However, these drugs can cause severe haemolysis in individuals with glucose-6-phosphate dehydrogenase (G6PD) deficiency. G6PD deficiency is caused by more than 230 different variants at the gene level that confer different degrees of deficiency phenotypically. Understanding the distribution of different G6PD variants in affected populations is essential to inform safer antimalarial treatment strategies. This study aimed to develop a cost-effective sequencing assay targeting key regions of the G6PD gene, suitable for field deployment. A novel assay based on Nanopore technology was designed to amplify two amplicons covering exon 3 to exon 13, focusing on known variants associated with enzyme deficiency. A total of 79 samples from individuals in Cambodia, Vietnam, Afghanistan, and China were sequenced, and a bioinformatics pipeline was created for the targeted variant calling of 192 G6PD SNP mutations. The assay demonstrated reliable detection of known variants, with high concordance between runs, within runs, and with Sanger sequencing.

The Nanopore MinION long-amplicon sequencing assay offers a robust and portable solution for large-scale G6PD genotyping in low-resource settings, that will improve malaria control and elimination strategies by enabling safer antimalarial treatment.

## Introduction

*Plasmodium vivax* remains an important public health threat in 41 endemic countries, causing 4–7 million cases of malaria each year [1]. Over the past two decades, intense malaria control efforts have reduced the global burden of disease considerably; however, in many areas where *P. vivax* and *P. falciparum* are co-endemic, *P. vivax* has become the predominant cause of malaria [1-3]. The primary reason for this shift in burden is the ability of *P. vivax* to causes recurrent episodes of malaria known as relapses. *P. vivax* and *P. ovale* are the only malaria parasites in humans capable of forming dormant liver stages (hypnozoites), which can reactivate weeks to months after an initial infection, causing recurrent parasitaemia. Up to 85% of recurrent *P. vivax* infections are attributable to relapses, and these contribute to significant morbidity and ongoing transmission of the parasite, which undermines elimination efforts [4,5]. Antimalarial treatment of *P. vivax* and *P. ovale* requires administration of a combination of schizonticidal and hypnozoitocidal drugs to kill both the asexual parasites, that cause acute disease, and the liver stages that result in relapses; this combination is referred to as ‘radical cure’. The only licensed drugs with hypnozoitocidal activity are primaquine and tafenoquine. Whilst these 8-aminoquinoline compounds are well tolerated in most patients, they can cause severe haemolysis in individuals with glucose-6-phosphate dehydrogenase (G6PD) deficiency (G6PDd) [6]. G6PDd is among the most common inherited enzymopathies worldwide, present in up to 35% of people living in malaria-endemic countries [7].

The *G6PD* gene has a size of ~18.5 kilobases (kb) and is located on the X chromosome (Xq28). It consists of 13 exons (the first exon is non-coding) and 12 introns, encoding a product of 1,545 base pairs (bp). Over 230 variants have been associated with reduced enzyme activity [8]. Most of these variants are single nucleotide polymorphisms (SNPs) in exon 3-13 [6,8,9]. Whilst males are either hemizygous-deficient or normal (wild-type), females can exhibit a more varied level of enzyme activity owing to the presence of two X chromosomes, one of which is randomly inactivated at the cellular level during early embryonic development through a process called lyonisation [6]. The degree of drug-induced haemolysis is related to the dose of drug administered and the underlying G6PD enzyme activity; the latter varies with the *G6PD* variant and the degree of lyonisation. In view of the risk of drug-induced haemolysis, the World Health Organization (WHO) recommends routine testing for G6PDd prior to treatment with either primaquine or tafenoquine. However, in practice, routine G6PD testing, despite the recent WHO pre-qualification of the SD Biosensor G6PD test, is rarely available in poor-resourced communities, where the greatest burden of *P. vivax* resides [10]. Geospatial mapping of different *G6PD* variants and their prevalence in *P. vivax* endemic countries can inform policy makers of the inherent risks of haemolysis if G6PD testing is not available, however such efforts have been confounded by the diversity of genotyping methods used [11].

Molecular methods such as polymerase chain reaction (PCR) followed by Sanger sequencing have been used to identify *G6PD* polymorphisms associated with enzyme deficiency. However, the application of these methods is usually tailored to identify variants known to be locally prevalent. While Sanger sequencing is accurate and useful for sequencing small genomic regions, it is less suitable for longer fragments due to its relatively high cost, slow processing time, and short read length compared to modern next-generation sequencing (NGS) technologies [12]. NGS platforms like Illumina are high-throughput and cost-effective but typically limited to 600 bp fragment lengths. In contrast, Nanopore sequencing stands out as one of the most advanced third-generation sequencing technologies, capable of generating sequence reads of varying lengths, from 20 kb to over 4 Mb [13], enabling detection of variants across the entire gene with fewer amplicons. The MinION sequencer from Oxford Nanopore Technologies is compact, portable, and relatively inexpensive compared to other sequencing platforms.

The aim of this study was to develop a novel MinION long amplicon-based sequencing assay spanning critical regions of the *G6PD* gene designed for deployment in resource-limited settings. Specifically, we sequenced 10.8 kb of the *G6PD* gene in two amplicons, covering from exons 3 to 13 and their flanking intronic regions. We validated the assay by confirming the presence of known *G6PD* mutations in patients from Cambodia, Afghanistan, China and Vietnam, using Sanger sequencing. To streamline data analysis, we developed an open-access bioinformatics pipeline to identify the 192 *G6PD* SNP variants associated with reduced enzyme activity, as listed by Luzzatto et al. [6]. Finally, we estimated the economic advantage of using the MinION platform for sequencing the *G6PD* gene in 96 samples as compared to Sanger sequencing by breaking down the costs associated with each method taking into account both the per-sample costs and the overall cost of sequencing.

## Materials and methods

### Sample collection and selection

Blood samples were collected from 79 individuals (43 females and 36 males) enrolled in cross-sectional surveys or antimalarial clinical trials conducted in Cambodia, Vietnam and Afghanistan, as well as a patient of Chinese ethnicity presenting at an Australian hospital (Supplementary Data 1). Individuals provided 250-400 µl of capillary blood or 4 ml of venous blood for screening of G6PD deficiency using either a qualitative test (Fluorescent Spot Test, FST) or a quantitative test (spectrophotometry or SD Biosensor). Female participants were categorized phenotypically as having G6PD normal enzyme activity (>70% of the adjusted male median activity, AMM), intermediate activity (30-70% of the AMM), or deficiency (<30% of the AMM). Males were categorized phenotypically as either normal (>30% of the AMM) or deficient (<30% of the AMM) [14]. Individuals tested with the FST were classified as G6PD-deficient if no fluorescence was observed after 30 minutes. For those tested using the SD Biosensor or spectrophotometry, G6PD activity below 4 IU/g Hb was classified as deficient, while activity between 4-6 IU/g Hb was considered intermediate [15]. The study was conducted following the declaration of Helsinki and all participants or their legal guardian provided informed consent.

### DNA extraction and Sanger sequencing

For 11 individuals enrolled in Afghanistan, China and Vietnam, DNA was extracted from whole blood using column kits (Favorgen Biotech, Taiwan). For 68 participants from Cambodia, DNA was extracted from whole blood using QIAamp DNA mini kit (Qiagen, Germany).

*G6PD* genotypes were determined by Sanger sequencing exon 3–13 following seven distinct PCR reactions using a protocol adapted from Kim et al. [16]. Chromatograms were visualized and analyzed with the Qiagen CLC Genomics Workbench (Qiagen, Germany). SNP mutations were identified using Mega-X software v.11 by comparison with the NCBI genomic RefSeqGene sequence NG_009015.2:g.

### Long-amplicon amplification, library preparation and MinION sequencing

The protocol, including primer sequences, PCR cycling conditions, and library preparation steps, can be found in Supplementary Information. Briefly, PCR amplifications were performed on two fragments of approximately 5.6 kb and 5.1 kb each, encompassing exons 3–13 and their flanking intronic regions (Figure 1). The reverse primer for the second fragment was positioned approximately 800 bp downstream of exon 13 to include its 3’ untranslated region. Amplification was performed using High Fidelity PrimeSTAR GXL DNA Polymerase (Takara Bio, Japan). The first fragment was obtained through two repeated PCRs utilizing the same primers. PCR products were run on 1% agarose gel for 1-1.5 hours at 100 V.

**Figure 1:**
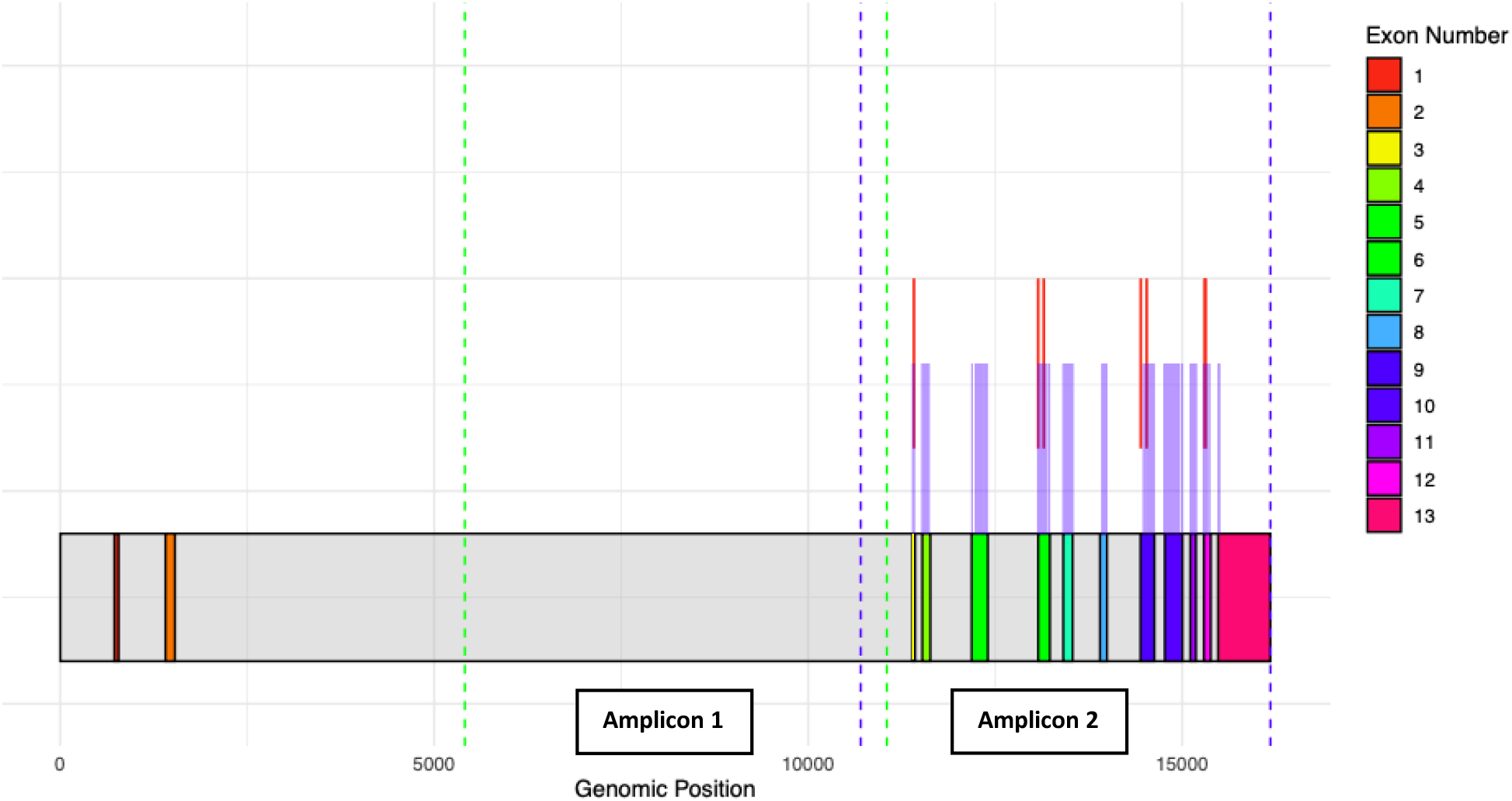
The *G6PD* gene, highlighting all exons from 1 to 13 (Transcript variant 3 from RefSeq NM_001360016.2:c, GenCode: ENST00000393562.10). The regions covered by Amplicon 1 and Amplicon 2 are indicated with green and dark purple dotted vertical lines, respectively. The position of variants that were identified in this study are marked with vertical red lines, while others included in the pipeline are shown in light purple.

A detailed list of the consumables and reagents used for the library preparation is provided by Oxford Nanopore at https://nanoporetech.com/document/native-barcoding-amplicons and summarised in Supplementary Information. Agencourt AMPure XP beads (Beckman Coulter, US) were used for high-throughput purification of PCR amplicons following the manufacturer’s instructions, maintaining a 1:1 ratio of beads to each PCR product during all purification steps. The Qubit Fluorometer 2.0 and Qubit dsDNA High Sensitivity Assay Kit (Thermo Fisher Scientific, US) were used to quantify PCR amplicons at each purification stage, tracking the DNA quantity to ensure it met the required levels for library preparation, including final pooling and loading onto the MinION flow cell.

Library preparation was carried out according to Oxford Nanopore Technologies instructions. Briefly, PCR amplicons of the two fragments from the same sample were quantified and pooled in equimolar amounts after purification, aiming for an ideal final concentration of 200-400 ng (120-240 fmol) per sample. DNA was then end-repaired, 5`phosphorylated, and prepared for ligation with the NEBNext Ultra II End Repair/dA-Tailing Module. To pool multiple samples onto one flow cell, the amplicons of each sample were labeled with unique barcodes using NEB Blunt/TA Ligase Master Mix. For the amplicon libraries of the first two runs performed (namely, run 1 and run 2) and the latter one (run 4), the ligation kit SQK-LSK109 with the amplicons-native barcoding expansion kit EXP-NBD104 was used, this was replaced by the expansion kit EXP-NBD196 for the 96-samples format of run 3. NEBNext Quick T4 DNA Ligase was used for the ligation of Nanopore adapters (Adapter Mix II) to the barcoded fragments, forming a complete library molecule. Before loading the prepared library, free oligonucleotides and small fragments were removed using AMPure XP beads. Lastly, priming of the flow cell, and loading of the DNA library was achieved using the flow cell priming kit (EXP-FLP002).

All runs were performed until exhaustion of the flow cell (up to 28 hours). For the pool libraries of run 1 and run 2, total reads were estimated at multiple times between 16 and 24 h for comparison in data yield. Since the coverage was highly satisfactory, when working in a 96-well plate, the clean-up step following end-prep was omitted to speed up the library preparation process, and an extended run time was included as a compensatory measure. Omitting the clean-up step and running until flow cell exhaustion allowed the initial volume and concentrations of pooled amplicons to be reduced (Supplementary Information). This method successfully processed final libraries with a concentration as low as 120 ng (below the recommended 200 ng) without any significant negative effects on sequencing quality, accuracy, or data yield. For run 2, ‘high accuracy basecalling’ was performed during the run while for the other runs including for multiplexing 96 samples, it was switched to ‘fast basecalling’ to improve computing time.

### Data analysis pipeline

A pipeline for targeted variant calling is available at https://github.com/vivaxgen/G6PD_MinION. A detailed breakdown of the pipeline steps can be found in Supplementary Information. After MinION sequencing, raw FAST5 reads were basecalled and demultiplexed using Guppy version 6.5.7 as a plugin in Genious Prime 2024. During the installation process, the reference sequence was automatically indexed (samtools version 1.20) [17]. The pipeline analysed the base-called sequencing data in five steps: i) trimming (chopper version 0.8.0) [18], ii) mapping (minimap2 version 2.28-r1209) [19], iii) variant calling (freebayes version 1.3.6 or clair3 version 1.0.9) [20,21], and, iv) a Python script that generates a variant report after applying quality-based filtering to remove low-confidence variant calls. Specifically, variants were filtered based on quality metrics (*e*.*g*., QUAL score) produced during variant calling, and empirical thresholds applied within the pipeline to minimize background noise while balancing sensitivity and specificity. These thresholds are not fixed globally, but rather are adapted based on sample characteristics, and are consistent with practices established in previous Nanopore-based variant calling workflows [22]. For the variant calling the pipeline was designed to take a list of known variants, to report the presence/absence of the variant as well as to characterize hemi-, homo-, and heterozygosity. The variants currently included can be found in Supplementary Data 2. The pipeline also has a variant discovery mode, which is still under development; this mode applies similar quality filtering criteria to candidate novel variants, but additional post hoc curation and validation steps are needed to mitigate the effects of noise and background.

## Results

### Assay development and samples sequenced

A total of 79 individuals from four different countries were sequenced in four different runs (Figure 2). In run 1, 3 samples were investigated, and ~7,000 reads per sample were obtained after 16 hours of sequencing, with an increase of ~6,000 passed reads at the 24-hour time-point. Similar numbers were observed when working with an increased number of samples, with over 10,000 reads per amplicon reached at 20 hours in run 2 (11 samples, data not shown). In run 3, to optimize MinION library preparation in a 96-well plate format and assess the impact of protocol modifications, samples from 23 individuals (2 females and 21 males) with known G6PD genotypes were sequenced in duplicates within the plate. Of these, one sample of Cambodian origin could not achieve concordant results within the run duplicate due to low read depth. Finally, run 4 (8 samples) was performed to provide a final check of the developed protocol. A total of 16 samples were run at multiple times across runs to assess reproducibility. All remaining samples were concordant between and within runs and showed agreement with Sanger sequencing results (Supplementary Data 1).

**Figure 2:**
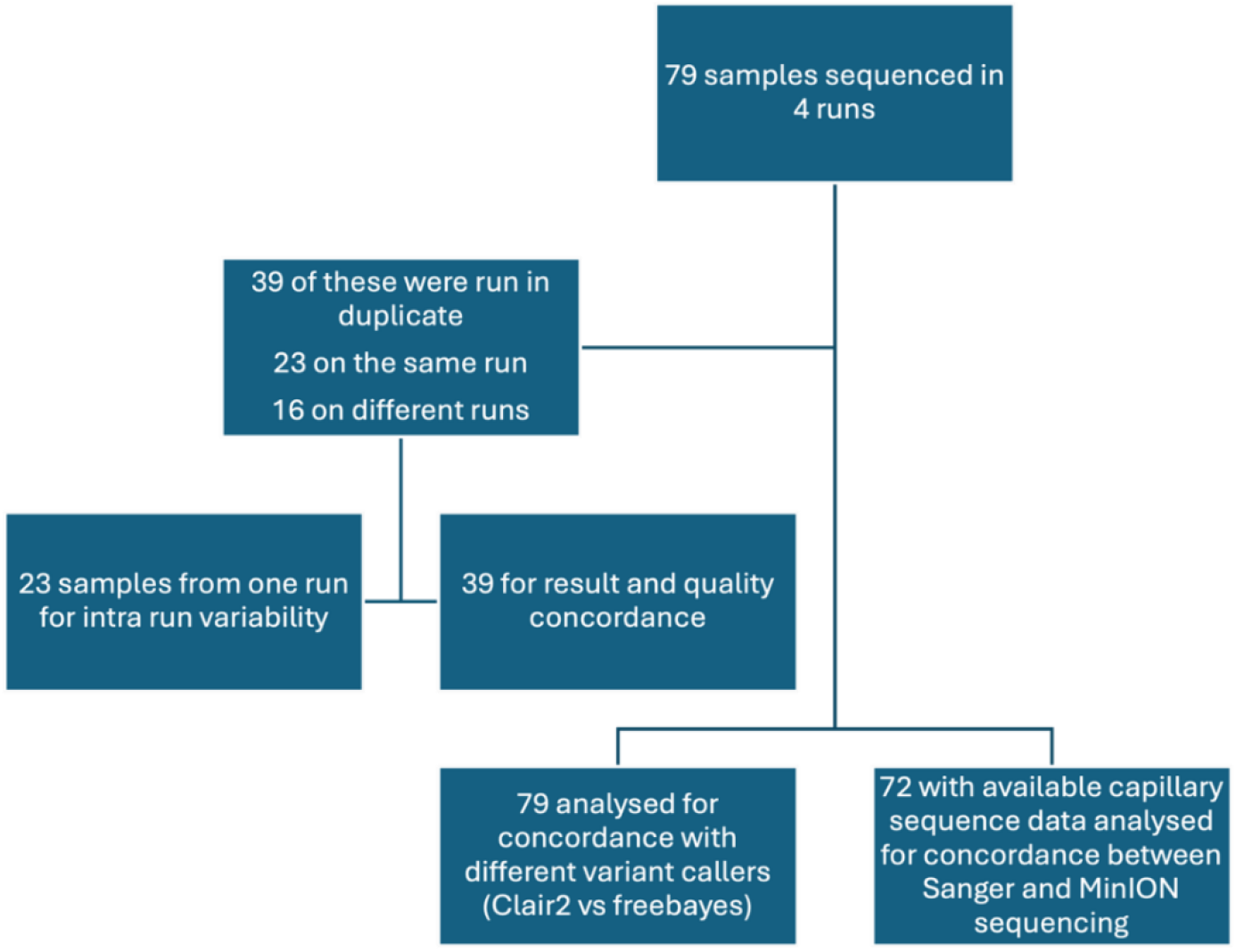
Overview of the samples selected for sequencing and their corresponding analyses.

### Sequencing performance

A summary of the data acquired at each run is given in Table 1. The mean number of reads per amplicon per sample varied from 686 to 94,641, resulting in a depth variation from 498 to 52,778 (Supplementary Data 3). These results were influenced by factors including running time, sample input concentration, available sequencing pores and sample multiplexing. For example, in run 2, an ultra-high depth of 52,000 was observed. Despite this variability, coverage and mapping quality were consistently high across the four runs, ranging from 97 to 100% for coverage and 59–60% for mapping quality. In the sequencing run with 96 samples, 2.77 million reads and approximately 5.14 Gb of bases were generated. Following analysis, the average depth was of ~950 for Amplicon 1 and of ~500 for Amplicon 2 (Figure 3).

**Table 1:**
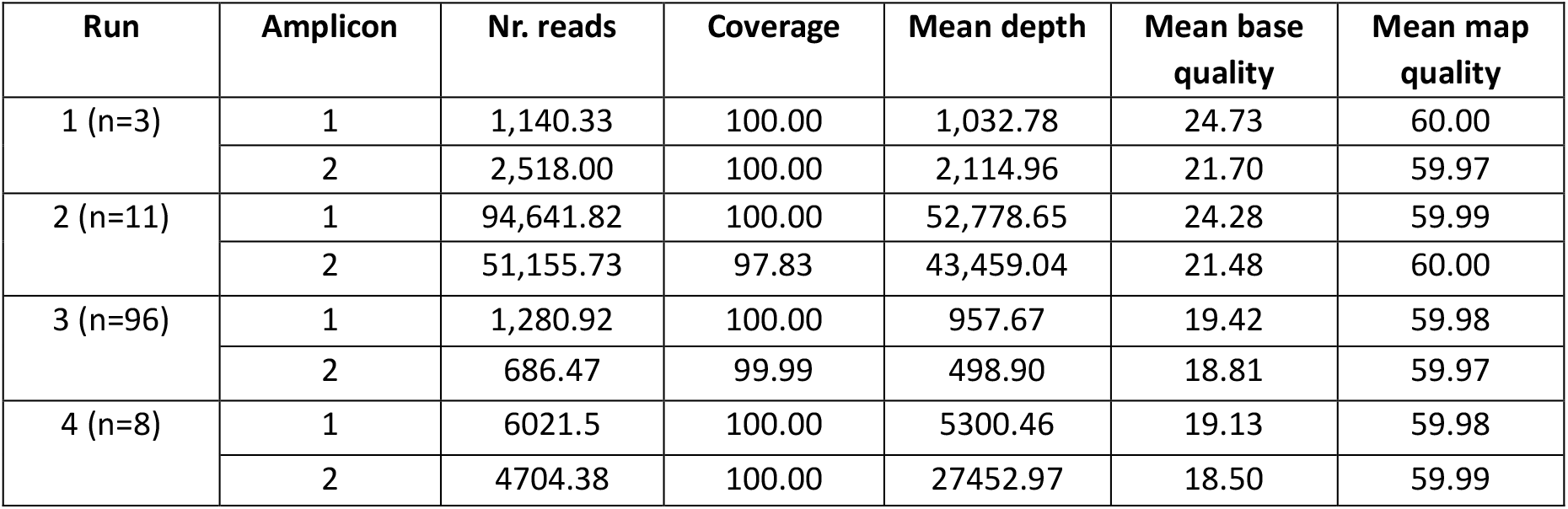
Summary of sequencing metrics per run and amplicon, including the average number of reads, mean depth, mean base quality, mapping quality and coverage. ‘n’ represents the number of samples processed at each run. All data shown were collected at flow cell exhaustion. Detailed metrics for each sample and run are available in Supplementary Data 3.

**Figure 3:**
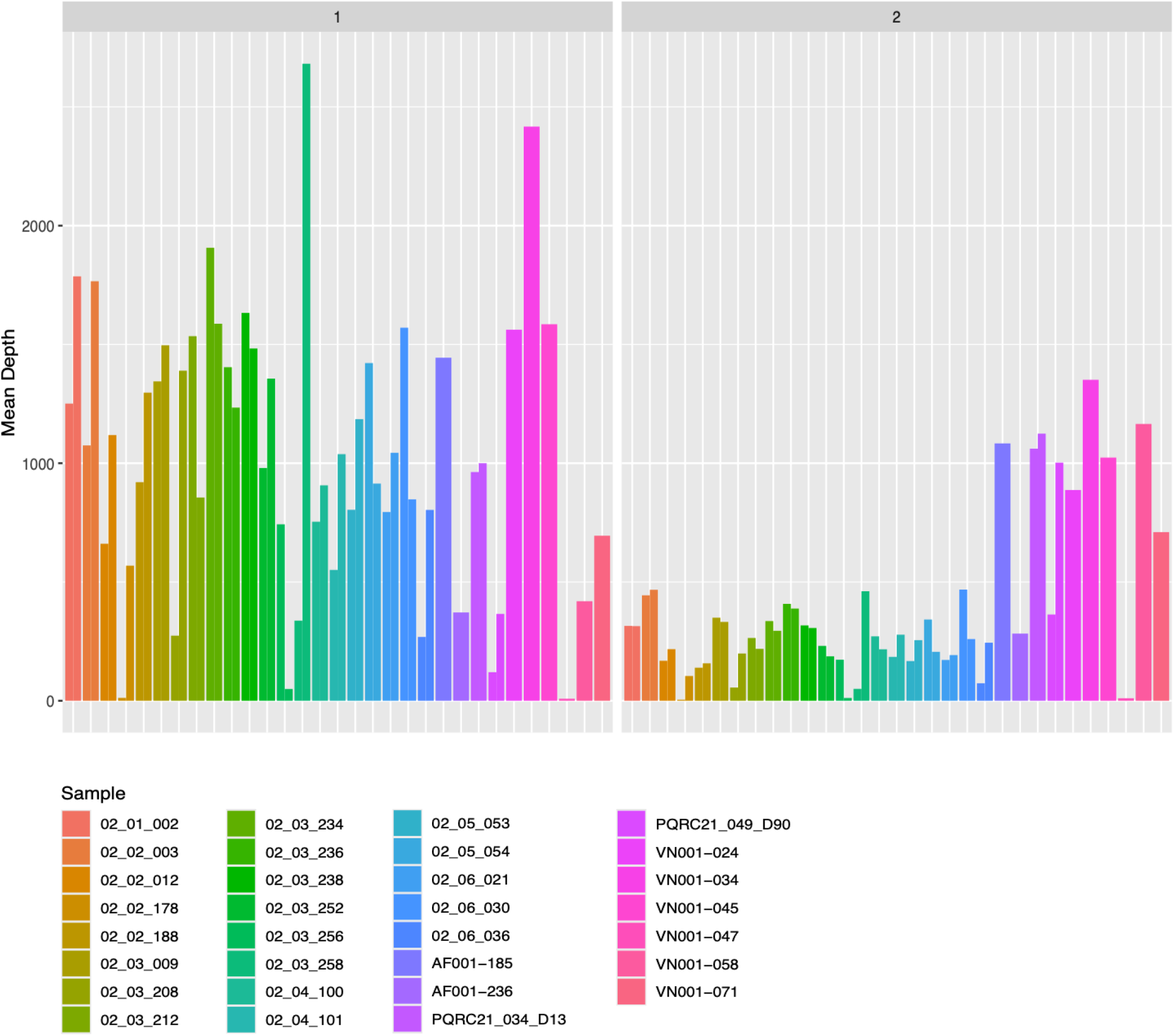
Mean sequencing depth for each sample. Each sample is represented by two bars of the same color, indicating duplicate sequencing runs associated with different barcodes. The left panel depicts the depth for Amplicon 1, while the right panel displays the depth for Amplicon 2.

### Assay validation and SNPs detection

The MinION pipeline was tested on samples from all 79 individuals, identifying seven known G6PD variants in 58 (73.4%). According to MinION results, of the 36 males included in the study, 58.3% (21/36) were hemizygous for G6PD variants, while the remaining 15 (41.7%) had no major known G6PD polymorphisms. Out of the 43 females sequenced, five (5/43, 11.6%) were homozygous for the G6PD Viangchan variant (Val291Met), and 28 (65.1%) were identified as heterozygous. Of these 28 heterozygous females, 26 carried the Viangchan variant, 1 had both the Viangchan and the Aureus (9Ile48Thr) variants, and 1 had the Mahidol (Gly163Ser).

Sanger sequencing was performed for 53 individuals carrying mutations according to MinION results (Table 2), and for 19 wild-type individuals to confirm absence of known polymorphisms. A final subset of 72 individuals had data from both Sanger and MinION sequencing: 61 from Cambodia, 3 from Afghanistan, 7 from Vietnam and, 1 from China (Supplementary Data 1). Five samples of Cambodian origins (2 males and 3 females) were not sequenced by Sanger due to low quantity of extracted DNA. Among the individuals carrying G6PD mutations, 43 (43/53, 81.1%) had exclusively Viangchan, 1 (1.9%) carried both Viangchan and Aureus variants, 3 (5.7%) had the Mediterranean variant (Ser188Phe), 3 (5.7%) had the Kaiping (Arg463His) and 1 each (1.9%) had the Canton (Arg459Leu), the Kerala-Kalyan (Glu317Lys), and the Mahidol variants. Complete concordance between Sanger and MinION was observed for all except one sample (Supplementary Data 1).

**Table 2:** Seven G6PD variants identified in 53 sequenced samples based on MinION results confirmed by Sanger sequencing. The table provides detailed information on each variant, including the corresponding ClinVar ID, and ClinVar interpretation. The reference sequences used were NCBI: NG_009015.2:g for genomic mutations, NM_001360016.2:c for mRNA mutations, and NP_001346945.1p for protein mutations (Isoform B).

| Name | ClinVar ID | ClinVar Interpretation | Genomic mutation | mRNA mutation | Protein change | Molecular Consequence | Exon |
| --- | --- | --- | --- | --- | --- | --- | --- |
| Aureus | 10402 | Pathogenic/Likely pathogenic | 16417T>C | 143T>C | p.Ile48Thr | Missense | 3 |
| Mahidol | 10367 | Pathogenic/Likely pathogenic | 18078G>A | 487G>A | Gly163Ser | Missense | 6 |
| Mediterranean | 100057 | Pathogenic | 18154C>T | 563C>T | Ser188Phe | Missense | 6 |
| Viangchan | 10386 | Pathogenic/Likely Pathogenic | 19451G>A | 871G>A | Val291Met | Missense | 9 |
| Kerala-Kalyan | 10401 | Pathogenic/Likely Pathogenic | 19529G>A | 949G>A | Glu317Lys | Missense | 9 |
| Canton | 100058 | Pathogenic | 20304G>T | 1376G>T | Arg459Leu | Missense | 12 |
| Kaiping | 100059 | Pathogenic | 20316G>A | 1388G>A | Arg463His | Missense | 12 |

### Cost comparison of Sanger and MinION sequencing of the *G6PD* gene

To highlight the potential cost savings that could be achieved through the adoption of the MinION platform, we evaluated the cost of reagents and consumables (including MinION flow cells and sequencer) required to sequence the *G6PD* gene for 96 and for 960 samples (Table 3).

**Table 3:**
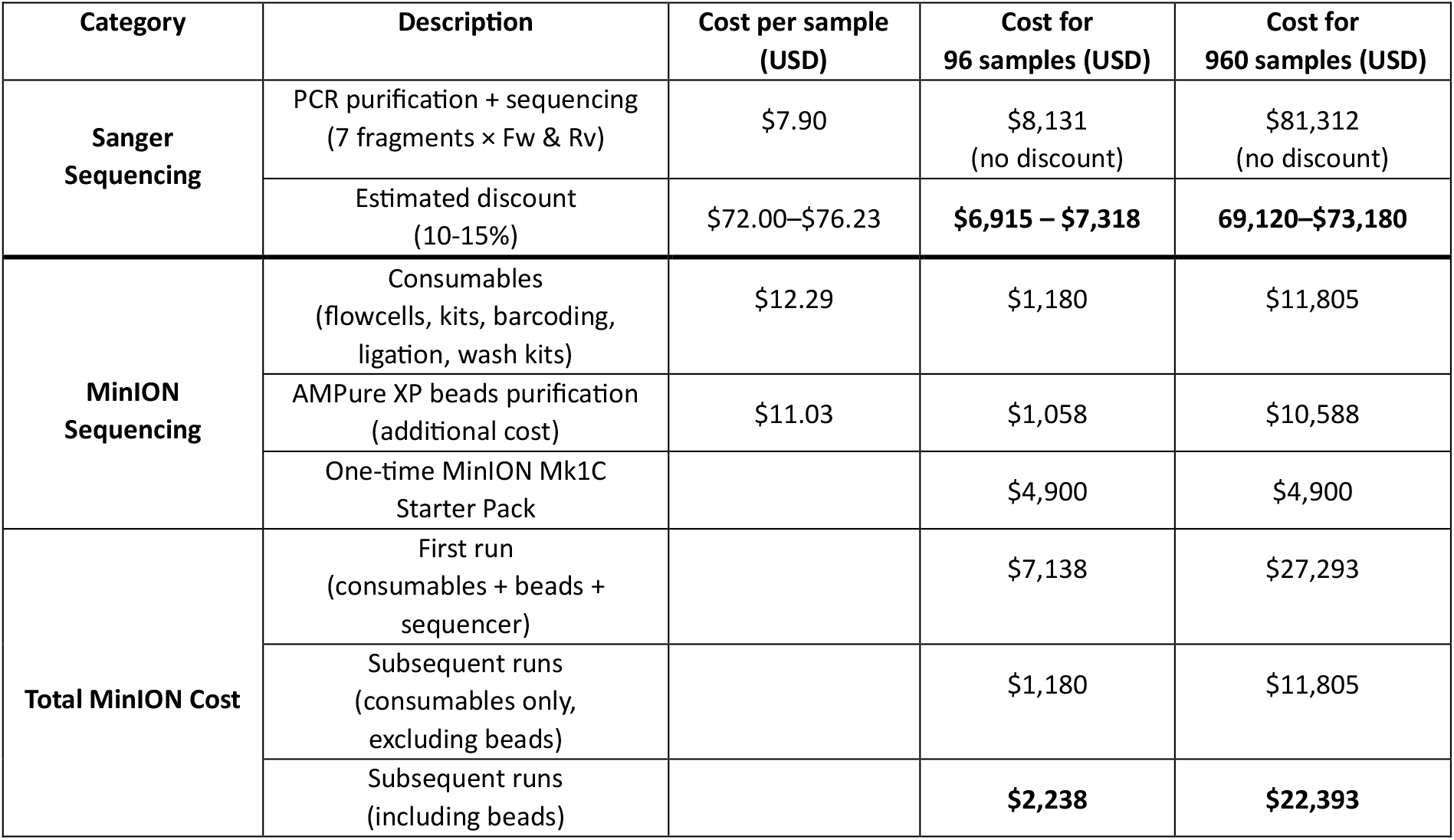
Cost comparison of Sanger and MinION sequencing for 96 and 960 samples of the *G6PD* gene. For Sanger, the price per sample includes seven PCR product purifications and 14 sequencing reactions. MinION consumables include flow cells, barcoding/adapters, end-prep/ligation enzymes, and wash kits, while AMPure XP beads are listed separately. The total MinION cost also includes the one-time purchase of the Mk1C sequencer.

| Category | Description | Cost per sample (USD) | Cost for 96 samples (USD) | Cost for 960 samples (USD) |
| --- | --- | --- | --- | --- |
| Sanger Sequencing | PCR purification + sequencing (7 fragments × Fw & Rv) | \$7.90 | \$8,131 (no discount) | \$81,312 (no discount) |
| | Estimated discount (10-15%) | \$72.00–\$76.23 | **\$6,915 – \$7,318** | **69,120–\$73,180** |
| MinION Sequencing | Consumables (flowcells, kits, barcoding, ligation, wash kits) | \$12.29 | \$1,180 | \$11,805 |
| | AMPure XP beads purification (additional cost) | \$11.03 | \$1,058 | \$10,588 |
| | One-time MinION Mk1C Starter Pack | | \$4,900 | \$4,900 |
| Total MinION Cost | First run (consumables + beads + sequencer) | | \$7,138 | \$27,293 |
| | Subsequent runs (consumables only, excluding beads) | | \$1,180 | \$11,805 |
| | Subsequent runs (including beads) | | **\$2,238** | **\$22,393** |

For Sanger sequencing, the entire coding exon regions is amplified using seven separate PCR reactions per sample, as described elsewhere [16,23]. Each of these PCR products was sent for purification and sequencing at the cost of $2.20 per purification and $4.95 per sequencing reaction (forward and reverse) to Macrogen, Korea, totaling $7.90 per PCR product. Therefore, the total cost for sequencing a single sample using Sanger was of $84.70 (7 purifications × $2.20 + 14 sequencing reactions × $4.95). However, with an estimated 10–15% discount commonly offered by sequencing providers, the Sanger sequencing cost per sample would decrease to $76.23–$72.00, respectively. For 96 samples, this would have amounted to $8,131 without discount, or $6,915–$7,318 after a 10–15% discount, and for 960 samples, the cost would scale linearly to $81,312 without discount or $69,150–$70,272 after discount.

In comparison, the MinION Mk1C Starter Pack was $4,900 and included a few kits, six flow cells and the MinION sequencer. Once purchased, the consumable costs for 96 samples (including flow cells, barcoding and adapter kits, ligation and end-prep enzymes, and wash kits), was about $1,180, that is, $12.29 per sample. The cost of AMPure XP beads (Beckman Coulter, US) purification was approximately $11.03 per sample ($1,058 total for 96 samples). Thus, the total MinION cost including beads was approximately $7,138 for 96 samples (sequencer + consumables + beads), or $2,238 for consumables and beads alone (no sequencer). For 960 samples (10 plates), consumables and beads alone would therefore cost approximately $22,380, and including the one-time sequencer purchase the total would be about $27,280. Even with the slightly higher price of the new MinION Mk1D Starter Pack ($4,950 including five flow cells), the cost advantage over Sanger sequencing remains substantial, especially as sample numbers grow.

Finally, in terms of turnaround time, MinION library preparation is done within a workday and results are delivered within 2-3 days for 96 samples on a same flow cell. On the other hand, PCR of 96 samples can be done in a single workday, as can be Sanger sequencing. However, as most laboratories do not have Sanger sequencing capacity which is quite expensive, PCR products have to be shipped to a sequencing centre, whether a commercial provider or academic collaborator, before being analysed and this can take weeks in malaria endemic areas.

## Discussion

We describe a novel sequencing assay and bioinformatics pipeline that reliably detects seven known G6PD variants associated with deficient enzyme activity, with the potential to identify up to 192 known variants (Supplementary Data 2). The MinION assay accurately genotyped 72 samples, as confirmed by Sanger sequencing. For these samples, we observed high concordance in sequencing results, even when a sample was run multiple times or in duplicate within the same run (Supplementary Data 1). Variant calls were also highly concordant when using two different callers, clair3 and freebayes (Supplementary Data 4) for both MinION and Sanger sequencing data, suggesting that our pipeline is robust across different bioinformatics tools. The only two discordant samples between runs identified during the study were obtained from two Cambodian male and female participants, for which Sanger sequencing was not performed (Supplementary Data 1). Of these, one sample (02_02_178) was run in duplicate during run 3 and was identified as homozygous for the Viangchan variant in one replicate and as wild-type in its within-plate duplicate, but the wild-type duplicate had very low read depth. The second sample, (02_02_092) was run twice (in run 3 and run 4). While in run 3 it showed low read depth consistent with a wild-type genotype, in run 4 it gave discordant results according to the two variant callers. Specifically, it was identified as heterozygous for the Viangchan variant by clair 3 and as homozygous for the same variant by freebayes. This discrepancy may have arisen from a variety of factors, including differences in variant calling thresholds. Variations in filtering parameters could influence allele frequency detection, potentially leading to false positive homozygous calls or missed heterozygous variants. Further investigation is needed to refine filtering criteria and improve the accuracy of genotype classification.

Sequencing depth varied between samples and runs, and was dependent upon the sequencing running time, fragments size and/or number of samples multiplexed. Sequence depth also varied for duplicates, and while DNA concentrations were quantified and equimolarly pooled at each run, the cause of this variability is unclear. Despite the depth variation, coverage and mapping quality remained consistently high across all runs, with coverage exceeding 97% and average mapping quality scores between 59 and 60. However, this variation may explain why a sample from a Cambodian female was identified differently as homozygous or heterozygous by the two sequencing methods.

The MinION assay identified seven known variants, with one of them having two different heterozygous variants. A total of 19 samples had no known variants detected according to both sequencing methods, of which 3 samples came from female individuals with intermediate G6PD activity (although one of these was close to normal with 68% of AMM activity) and 4 from males phenotypically diagnosed as deficient (Supplementary Data 1). Of these, 1 had G6PD activity determined by both spectrophotometry and by SD Biosensor, and the two tests led discrepant results, while for the other 3 enzyme activity was assessed using the FST method. This could indicate the influence on enzyme activity of novel variants that have yet to be described in the literature or an error associated to the testing. Further work is now underway to adapt the pipeline for variant discovery and laboratory confirmation.

Geographical mapping of the prevalence of different G6PD variants is complicated by the variety of methods used in different surveys and countries [11]. In our Cambodian dataset, all 68 samples were collected from individuals in Kampong Speu province, located in Western Cambodia. MinION sequencing identified the Viangchan mutation as the most common variant, present in 47 samples (69.1%) (Supplementary Data 1). In contrast, samples from Afghanistan carried the Mediterranean variant, while samples from Vietnam showed Canton, Kaiping and Viangchan mutations. However, the small sample sizes and non-representative sampling in these countries limit the ability to draw broad conclusions about variant distribution and prevalence at population level.

Our study has several limitations. The assay was designed to detect only variants within exon 3–13 excluding the gene promoter region, the first non-coding exon, and exon 2. Additionally, the assay was also assessed in a limited number of geographical areas and known variants, using venous blood samples, so further investigation of other populations and sampling types, such as dried blood spots, is warranted. Nevertheless, the application of this standardized assay in larger populations has significant potential to enhance our understanding of *G6PD* genetic variants in *P. vivax*-endemic regions, including association of different variants with enzyme deficiency and associated haemolytic risk in patients treated with primaquine or tafenoquine. The assays will also generate valuable insights into *G6PD* mutations in both coding and non-coding regions of the gene, improving our understanding of genotype-phenotype correlations and their implications for the diagnostics of G6PD deficiency.

In terms of cost-effectiveness, MinION sequencing offers a far more economical approach for sequencing *G6PD* gene than traditional Sanger sequencing given 96 samples per flow cells are run at a time, emphasizing its suitability for large-scale studies. Additionally, MinION significantly reduced turnaround time, with results delivered within 2–3 days, whereas Sanger sequencing can take weeks depending on the sequencing provider’s location.

## Conclusions

The choice of sequencing technology for gene analysis can significantly impact both the financial and logistical aspects of a study, especially when dealing with large sample sizes. We present a novel long-amplicon sequencing assay that reliably detected *G6PD* mutations in exons 3-13 in Asian populations from Cambodia, Afghanistan, Vietnam and China. Our results demonstrate that MinION sequencing is a cost-effective and time-saving alternative to Sanger sequencing. Additionally, this study highlights the reliability of MinION sequencing, making it a recommended choice for *G6PD* genotyping in large-scale studies such as cross-sectional surveys and clinical trials. These advantages are particularly valuable in resource-limited settings, where efficiency and affordability are essential.

## Data availability

The data supporting the findings of this study are available in the European Nucleotide Archive (ENA) under the following primary accession number: PRJEB88995

## Code availability

The pipeline for obtaining SNP data in Variant Call Format (VCF) is accessible on GitHub at: https://github.com/vivaxgen/G6PD_MinION.

## Ethical approval

Ethics approval was obtained from the following national and local committees and authorities: The Human Research Ethics Committee of the Northern Territory Department of Health (HREC), Australia, the Islamic Republic of Afghanistan, Ministry of Public Health, Institutional Review Board, Afghanistan, the Oxford Tropical Research Ethics Committee (OxTREC), UK, the National Ethics Committee for Health Research, Cambodia, the U.S. Centers for Disease Control and Prevention, and the Ministry of Health Evaluation Committee on Ethics in Biomedical Research, Vietnam.

## Funding

The collection of patient samples was funded by the U.S. President’s Malaria Initiative, USA, the UK Department for International Development, UK Medical Research Council, UK National Institute for Health Research, and the Wellcome Trust through the Joint Global Health Trials Scheme (MR/K007424/1) and the Bill & Melinda Gates Foundation [INV-024389]. M.N. was supported by the IDEX International Master’s Scholarship from Université Paris-Saclay, France. R.N.P. and S.A. are funded by an Australian National Health and Medical Research (NHMRC) Leadership Investigator Grants (2008501 and 2001083). J.P. is supported by the NIH/NIAID (R01AI173171, R01AI175134 and R61AI187100). J.P. and BW are supported by the Pasteur International Unit PvESMEE.

## Disclaimer

The findings and conclusions in this report are those of the author(s) and do not necessarily represent the official position of the Centres for Disease Control and Prevention or the United States Agency for International Development.

## Author contributions

I.M., R.N.P. and J.P. conceived the study. C.T. and M.K. designed the study. C.T. designed the laboratory assay. C.T., R.E., M.N., N.K. and A.R. contributed to sequencing data generation and laboratory data analysis. C.T., M.K., H.T., K.S.H., and M.N. developed the bioinformatic pipeline and conducted the bioinformatic data analysis. B.L., J.H., B.W., S.A., I.M., R.N.P. and J.P. contributed to essential field-based sample collections and metadata, and guidance on the study design and interpretation. C.T., M.K., S.A., R.N.P. and J.P. wrote the manuscript. All authors approved the final version of the manuscript.

## Notes

### Competing Interest Statement

The authors have declared no competing interest.

