## Supplementary material for "Rapid detection of G6PD deficiency SNPs using a novel amplicon-based MinION Sequencing Assay": Supp text

*equal contribution

Corresponding author: Jean Popovici

### **Supplementary Table 1.** Primer Sequences used for *G6PD* sequencing of Amplicon 1 and Amplicon 2. The nucleotide start and end position are referenced to the NCBI genomic RefSeqGene sequence NG_009015.2:g.

| **Amplicon Name** | **Primer name** | **Primer Description** | **Primer Sequence (5’-3’)** | **Nucleotide Position** | **Length**  **(bp)** |
| --- | --- | --- | --- | --- | --- |
| Amplicon  1 | G6PD-Fw1 | Forward | CCCATTGGTCTGATAAGATTCTTGC | 10,413 - 10,438 | 5,643 |
|  | G6PD-Rv1 | Reverse | AGGAGCTAAGCATTCCACTTAAGAA | 16,031 - 16,056 |  |
| Amplicon  2 | G6PD-Fw2 | Forward | TGGCTTGCAGTTCTATCTTTTTC | 16,349- 16,368 | 5,135 |
|  | G6PD-Rv2 | Reverse | ATACTTCTGTGGACTGGCAGTGT | 21,461- 21, 484 |  |

### **Supplementary Table 2.** PCR conditions and program for the amplification of Amplicon 1 and Amplicon 2. For Amplicon 1, a nested PCR was performed using the same primers, reagents, and program conditions as the primary PCR, with two key differences: 0.5 µL of the primary PCR product was added to the master mix, and the denaturation-annealing-extension cycle was repeated 25 times instead of 30. Amplicon 2 was successfully amplified with optimal yield without requiring a nested PCR. All PCR products were run on 1% agarose gel at 100 V for 1 -1.5 hours.

**Amplicon 1**

PCR conditions for primary PCR

|  | **Reagent** | **Stock Conc.** | **Final Conc.** | **Vol. for one sample** |
| --- | --- | --- | --- | --- |
| 1. | Sterile purified water |  |  | 31.5 μl |
| 2. | PrimeSTAR Buffer | 5X | 1X | 10 μl |
| 3. | dNTP Mixture | 2.5 mM | 0.2 mM | 4 μl |
| 4. | Forward primer | 10 μM | 0.25 μM | 1.25 μl |
| 5. | Reverse primer | 10 μM | 0.25 μM | 1.25 μl |
| 6. | PrimeSTAR® GXL DNA Polymerase | 2.5 U/μl | 1.25 U | 1 μl |
|  | gDNA | >50ng/ul | >1ng/ul | 1ul |
|  | *Total volume of reaction* |  |  | *50 ul* |

PCR program for primary PCR (Expected time: ~ 4 hours)

| **Step no.** | **Cycle** | **Temperature (ºC)** | **Time (min)** | **No. of cycles** |
| --- | --- | --- | --- | --- |
| 1. | Preheating | 98 | 3:00 |  |
| 2. | Denaturation | 98 | 0:10 | 30x total |
| 3. | Annealing | 55 | 0:15 |  |
| 4. | Extension | 72 | 6:00 |  |
| 5. | Final extension | 72 | 5:00 |  |
| 6. | Pause | 4 | ∞ |  |

**Amplicon 2**

PCR conditions

|  | **Reagent** | **Stock Conc.** | **Final Conc.** | **Vol. for one sample** |
| --- | --- | --- | --- | --- |
| 1. | Sterile purified water |  |  | 31.5 μl |
| 2. | PrimeSTAR Buffer | 5X | 1X | 10 μl |
| 3. | dNTP Mixture | 2.5 mM | 0.2 mM | 4 μl |
| 4. | Forward primer | 10 μM | 0.25 μM | 1.25 μl |
| 5. | Reverse primer | 10 μM | 0.25 μM | 1.25 μl |
| 6. | PrimeSTAR® GXL DNA Polymerase | 2.5 U/μl | 1.25 U | 1 μl |
|  | gDNA | >50ng/ul | >1ng/ul | 1ul |
|  | *Total volume of reaction* |  |  | *50 ul* |

PCR program (Expected time: ~ 2 hours)

| **Step no.** | **Cycle** | **Temperature (ºC)** | **Time(min)** | **No. of cycles** |
| --- | --- | --- | --- | --- |
| 2. | Denaturation | 98 | 0:10 | 30x total |
| 3. | Annealing | 60 | 0:15 |  |
| 4. | Extension | 68 | 4:00 |  |
| 5. | Final extension | 68 | 0:15 |  |
| 6. | Pause | 4 | ∞ |  |

### **Supplementary Note 1.** List of kits and reagents required for library preparation from New England Biolabs® (NEB) and Oxford Nanopore Technologies® according to library preparation step.

1. End-prep

- NEBNext® UltrA II End Prep Enzyme Mix and
- NEBNext® Ultra II End Prep Reaction Buffer

from the NEBNext® Ultra II End repair/dA-tailing Module (NEB #E7546) or from the NEBNext® Companion Module for Oxford Nanopore Technologies® Ligation Sequencing (NEB #E7180S). The latter kit additionally contains a Quick T4 DNA Ligase for adapter ligation.

2. Native barcoding

- Native Barcoding Expansion kit 96 (ONT #EXP-NBD196)
- NEB Blunt/TA Ligase Master Mix (NEB #M0367)

3. Adapter ligation

- Long Fragment Buffer (LFB) from Ligation Sequencing kit (ONT #SQK-LSK-109)
- Elution Buffer EB from Ligation Sequencing kit (ONT #SQK-LSK-109)
- Adapter Mix (AMX) from Native barcoding expansion kit. It can be replaced by the Adapter Mix II (AMII) from Adapter Mix II Expansion (ONT #EXP-AMII00001)
- NEBNext® Quick Ligation Reaction Buffer (NEB #B6058) and
- NEBNext® Quick T4 DNA Ligase (NEB #E6057)

from NEBNext^®^ Companion Module for Oxford Nanopore Technologies^®^ Ligation Sequencing OR from NEBNext® Quick Ligation Module (#E6056).

4. Priming

- Flow Cell Priming kit (ONT #EXP-FLP002)
- Loading Beads (LB) from Ligation Sequencing kit (ONT #SQK-LSK-109)
- Sequencing Buffer (SQB) from Ligation Sequencing kit (ONT #SQK-LSK-109)

### **Supplementary Note 2.** MinION library preparation for long-amplicon sequencing of 96 samples. Samples used for step A were previously amplified by PCR, and the PCR products were quantified by Qubit™ Fluorometer 2.0 and Qubit™ dsDNA High Sensitivity Assay Kit, as per manufacturer’s instruction.

#### **A. Purification of PCR products**

PCR product purification for Amplicon 1 and Amplicon 2 was carried out using AmPure XP Beads, following the manufacturer's instructions with minor modifications (available at: https://www.beckman.it/reagents/genomic/cleanup-and-size-selection/pcr#WorkflowProtocol).

When multichannel pipettes, 96-well magnetic separation racks, and a compatible centrifuge are available, the initial purification step can be performed using two 96-well plates, one for each amplicon. However, since the first purification does not need to be done simultaneously for all samples, we recommend using Eppendorf DNA LoBind tubes, preparing one tube for each sample to be purified.

Note: Amplicon 1 and Amplicon 2 from the same sample should be processed separately, each in its own well or tube.

Briefly:

1. Allow the beads to thaw at room temperature (RT), and vortex them immediately before use to resuspend any settled magnetic particles. Add 40 µL of beads to each 40 µL PCR product, maintaining a 1:1 ratio, and mix by gently pipetting 10 times. The mixture should appear homogenous after mixing. If the amplicon concentration exceeds 50 ng/µL (ideally over 100 ng/µL), less PCR product can be used—down to as little as 10 µL. However, a minimum of 20 µL is recommended to ensure sufficient product yield for pooling. It is important to always maintain a 1:1 ratio of beads to PCR product.
2. Incubate for 10 minutes at RT to allow DNA binding to the beads, preferably using a rotator mixer. Ensure the tube lids are properly closed to prevent contamination. During this incubation, freshly prepare 70% ethanol in nuclease-free water (~500 μl/product).
3. After incubation, place the tubes/plate onto a magnetic rack to separate the beads from the supernatant (approximately 2-5 minutes).
4. Once the solution is clear, discard the supernatant using either a multichannel pipette or a P200 pipette, taking care not to disturb the beads.
5. Add 200 µl of freshly prepared 70% ethanol to each well or tube while the plate or tubes remain on the magnetic rack. Note: Removing the supernatant and adding ethanol immediately will facilitate easier resuspension of the beads.
6. Remove the ethanol as described in step 4 and repeat the addition of 70% ethanol as outlined in step 5.
7. Perform a pulse-spin in a microfuge to remove any residual ethanol. Allow the pellet to air dry until it no longer appears shiny (approximately 30 seconds), being careful not to crack the beads.
8. Remove one tube at a time from the magnetic rack, add 30 µl of nuclease-free water to each tube, and mix by pipetting up and down at least 10 times. Visually inspect the solution to confirm that no clumps remain.
9. Pulse-spin the samples in a microfuge, place each tube back on the magnetic rack, and wait until the solution becomes clear. Prepare a new set of labeled 1.5 ml Eppendorf tubes for each final cleaned PCR product and arrange a basket with ice.
10. Transfer 28 µl of the eluted solution into the corresponding labeled clean tubes. Avoid aspirating any beads during the transfer. Ensure that the tubes are kept on ice throughout this process to prevent degradation. It is important to handle the eluted solutions carefully to avoid contamination and ensure accurate transfer. Once all transfers are complete, store the eluted products at 4°C, on ice or proceed directly to the next step in the protocol.
11. Quantify 1ul of purified PCR product by Qubit Fluorometer 2.0.

#### **B. Amplicon pooling, End Prep procedure and Native barcode ligation**

After purification, amplicons from the same sample are pooled in equimolar amounts. It is essential that all samples follow library preparation with the same initial concentration. It is important to identify the amplicon with the lowest concentration across all samples and adjust the concentrations of the other amplicons accordingly.

Recommended Amplicon concentration:

For optimal pooling, a minimum concentration of approximately 8.5 ng/µl per amplicon is recommended. This ensures a starting concentration of 200 ng per amplicon in a final volume of 24 µl, which corresponds to 400 ng per sample in the required 48 µl volume for End Prep. If necessary, this concentration can be reduced to 120 ng per amplicon (equivalent to 5 ng/µl).

In cases where PCR product quality and concentrations are satisfactory, the purification step following End Prep may be removed to optimize time. Consequently, the End Prep volume can be reduced by half, as only 22.5 µl of end-prepped DNA is required for barcoding ligation. This allows the use of 200 ng per amplicon in a final volume of 12 µl (yielding a concentration of 16.7 ng/µl).

Amplicons that cause a significant reduction in the overall concentration should be excluded from the library preparation and may require further amplification.

For the detailed End Prep procedure and native barcode ligation, please refer to Step 4 and Step 5 of the Oxford Nanopore protocol for native barcoding of amplicons, available at:

https://nanoporetech.com/document/native-barcoding-amplicons.

#### **C. Sample pooling**

Adapter ligation is performed on a single pooled sample, composed of all individual barcoded samples. The final required volume for adapter ligation is 65 µl with a total concentration as low as 150 ng. Therefore, the volume and concentration of each sample must be adjusted accordingly. Divide the total ligation volume by the number of pooled samples. Dilute each sample with nuclease-free water to match the least concentrated sample, ensuring uniformity.

#### **D. Adapter Ligation Protocol, final purification and elution of DNA library**

1. Thaw and prepare buffers:
   - Thaw Elution Buffer (EB) and NEBNext Quick Ligation Reaction Buffer (5X) at room temperature (RT). Mix by vortexing, spin down, and place on ice. Ensure that all reagents are clear of any precipitate.
   - Prepare Enzymes and Adapter Mix: spin down T4 Ligase and Adapter Mix II (AMII) and place on ice.
   - Thaw Long Fragment Buffer (LFB) at RT. Mix by vortexing, spin down, and place on ice.
2. Set up the ligation reaction (mix by flicking the tube between each addition):

| **Reagent** | **Volume** |
| --- | --- |
| Pooled barcoded DNA | 65 µl |
| Adapter Mix II (AMII) | 5 µl |
| NEBNext Quick Ligation Reaction Buffer (5X) | 20 µl |
| Quick T4 DNA Ligase | 10 µl |
| *Total Volume* | *100 µl* |

3. Mix thoroughly by pipetting and spin down. Incubate for 10 minutes at room temperature.
4. Purify the sample by adding 50 µl of resuspended AMPure XP beads (2:1 sample-to-bead ratio).
5. Mix thoroughly by pipetting and incubate on a rotator mixer for 5 minutes at RT.
6. Place on a magnetic rack, wait for beads to pellet, then carefully pipette off the supernatant.
7. Wash beads (repeat twice) as follow:
   - Add 250 µl Long Fragment Buffer (LFB) and flick to resuspend beads.
   - Spin down, return to the magnet, and remove supernatant.
8. Dry and resuspend:

- Spin down and place back on the magnet.
- Remove any remaining supernatant.
- Air dry for ~30 seconds, but do not over-dry (avoid pellet cracking).
- Remove the tube from the magnetic rack and resuspend in 15 µl Elution Buffer (EB).
- Spin down and incubate for 10 minutes at room temperature.

9. Place back on the magnet and wait at least 1 minute until the eluate is clear and colourless.
10. Recover DNA library: transfer 15 µl of eluate (containing the DNA library) into a clean 1.5 ml Eppendorf tube.

For the final steps of priming and loading the library molecule onto the MinION flow cell, please follow instruction as per Oxford Nanopore protocol for native barcoding of amplicons.

### **Supplementary Note 3. Bioinformatics Pipeline for targeted G6PD variant calling of MinION sequencing data**

Variant calling from fastq files using either freebayes or clair3 were incorporated into the data processing pipeline, which is available at <https://github.com/vivaxgen/G6PD_MinION>. The raw fastq files were filtered and trimmed with chopper. The trimmer trims 40 bases from both the start and end of each read. Additionally, the trimmer retains only reads that have a minimum 10 Phred average quality score, and is between 2,000 and 10,000 bases. The trimmed sequences were mapped to the NCBI reference: NC_000023.11 with minimap2 and the resulting bam file was sorted with samtools. Variant calling can be performed with either freebayes or clair3. When freebayes is used, heterozygous calls require a minimum of five reads supporting the alternate allele, with the alternate allele constituting at least 25% of the total read depth. For clair3, variants were performed with r941_prom_sup_g5014 model, with the estimated base error rate set at 0.05, and the minimum allele frequency supporting SNP and INDEL were both set at 0 while all other parameters were retained at default. The resulting VCFs were inspected with a python script that inspects each of the interested variants and generate a final report comprising of all the input samples and indicates whether any G6PD-causing variants were detected, and whether the detected variants were homo- or heterozygosity.
